# Upregulation of *GREB1* in colorectal cancer ovarian metastases may be a potential therapeutic target

**DOI:** 10.1101/2025.08.31.673371

**Authors:** Samantha M. Ruff, Alessandro La Ferlita, Manoj Palavalli, Joal Beane, Susan Tsai, Oliver S. Eng, Alex C. Kim, Ashwini K. Esnakula

## Abstract

**Introduction:** Classification of ovarian metastases (OM) in colorectal cancer (CRC) as peritoneal metastasis (PM) remains controversial. OM demonstrate resistance to systemic therapy, suggesting distinct molecular tumorigenesis. Gene expression profiling and transcriptomic analysis were performed to distinguish OM, PM, and primary CRC (pCRC) and identify potential therapeutic targets.

**Methods:** RNA sequencing data were obtained from the Total Cancer Care database. After filtering out low-expressed genes, raw counts of retained genes were normalized in trimmed mean of M-values and then log2-transformed. Genes with a |Log2FC|>0.58 and adjusted p-value <0.05 were considered dysregulated. In silico motif enrichment analysis of estrogen receptor 1 (ESR1) on growth regulating estrogen receptor binding 1 (GREB1) was conducted.

**Results:** There were 115 patients with tissue from OM (n=6), PM (n=4), and pCRC (n=105). Among upregulated genes, LAMC3, SCUBE1, and GREB1 had the highest differential expression in OM compared to PM, and PEG3, C7, and GREB1 had the highest differential expression in OM compared to pCRC (all p<0.001). GREB1 was upregulated in OM compared to PM and pCRC. There are two estrogen response element (ERE) sites within the promoter region of GREB1. ESR1 transcription factor was shown to bind to these ERE (p<0.001).

**Conclusions:** Transcriptomic analysis demonstrated clear molecular distinction between OM, PM and pCRC. We identified upregulation of GREB1 in OM compared to PM and pCRC. GREB1 contains both ERE and ESR1 in the upstream promoter regions implying potential upstream transcription regulation mediated by ESR1. GREB1 may represent a novel, hormonal target in treatment of OM in CRC.

## Introduction

Colorectal cancer (CRC) is the second leading cause of cancer death in the United States and represents an increasing healthcare burden.[1] About 25% of patients with colorectal cancer present with synchronous metastatic disease, while another 20% will develop metachronous metastases during their disease course.[2] As the second leading cause of cancer-related death in the world, it is critical that research remains focused on understanding the underlying mechanisms of metastatic spread, preventing the development of macrometastatic disease, and personalizing treatment for patients with metastatic CRC. Liver, lung, and peritoneum are the most common sites of metastatic disease, with peritoneal metastases (PM) associated with a worse prognosis.[3-6] PM are classified as any implant within the peritoneal cavity outside of the liver and retroperitoneum. This classification is controversial, particularly regarding ovarian metastases (OM) due to contrasting biological differences.[7] OM show resistance to systemic therapy, suggesting distinct molecular tumorigenesis from PM.

Metastatic sites (e.g., liver, peritoneum, ovary) exhibit different responses to the same systemic therapies and correlate with prognosis, suggesting biologic differences between them. The route of spread and unique native microenvironment of different metastatic sites likely selects for a specific population of circulating cancer cells. Cancer cells interact with the native tissue at different metastatic sites to establish an ideal tumor microenvironment (TME) that promotes the transition from micro to macrometastatic disease. A large proportion of research is focused on the circulating and resident immune cells, but not on native epithelial cells. Epithelial cells are abundant throughout the body, and their function changes based on anatomic location. The underlying mechanisms of how epithelial cells at distant metastatic sites interact with cancer cells to promote survival and proliferation are unknown. These interactions are difficult to study because pre-clinical models often do not include epithelial cells or consist of epithelial cells from animal models that are not representative of human cell behavior. Understanding the differences in gene expression between different metastatic sites (e.g. liver, lung, ovary, peritoneum) may help identify new directions for research and potential targets for systemic therapy. This is especially true for PM and OM, given that the underlying mechanism of metastatic spread is still unknown. This study set out to compare the gene expression profile of OM, PM, and primary CRC to identify potential therapeutic targets.

## Methods

### Data Source and RNA Sequencing Analysis

Clinical and molecular data were obtained from the Total Cancer Care (TCC) network. RNA sequencing data of OM (n=6), PM (n=4), and primary CRC (n=105) samples were obtained as not-normalized raw counts from the Oncology Research Information Exchange Network (ORIEN) TCC Program after data request approval. Raw counts were scaled using the Reads Per Million (RPM) formula to filter out low-expressed genes prior to normalization and differential expression analysis. Precisely, all the genes whose geometric mean of the RPM was less than one across all samples were removed. Afterward, raw counts of retained genes were normalized in trimmed mean of M-values (TMM) and then log2-transformed using the Limma R package.[8] Genes with a |Log2FC|>0.58 (|Linear FC|>1.5) and an adjusted p-value <0.05 (Benjamini-Hochberg correction) were considered differentially expressed. All the analyses have been performed in R (v. 4.2.2) using the RStudio (v. 2022.12.0) framework.

### In Silico Analysis of Transcription Factor Binding

Estrogen response elements (ERE) have been previously identified as a consensus palindromic sequence.[9] This sequence was searched 5KB to the start codon of GREB1, showing 2 specific ERE. ESR1(also known as ER-α) is the transcription factor for estrogen and specifically binds to these ERE.[10] Find Individual Motif Occurrences (FIMO) analysis of ESR1 5KB upstream of GREB1 was conducted with a cutoff p-value < 1e^-5^. These results were then cross-analyzed with ERE.

### Immunohistochemistry Staining for ESR1 and GREB1

This study was approved by the Institutional Review Board at The Ohio State University (IRB# 2024C0019). Immunohistochemical detection of GREB1 and ESR1/ER-α was accomplished using mouse monoclonal anti-GREB1 antibodies (MABS62, Millipore sigma, MO, USA) and Anti-SP1 rabbit polyclonal antibody clone for ESR1/ER-α (Abcam, MA, USA) with Leica/Bond polymer detection system or Agilent Envision Flex detection system, on a Leica/Bond (Il, USA) or Agilent DAKO autostainers (CA, USA). A dilution of 1:100 for Anti-GREB1 and 1:600 for Anti-SP1 antibodies was considered optimal after validating the staining protocols using appropriate tissue controls. The immunohistochemical stain slides were reviewed by a pathologist (AE) for expression of GREB1 and ESR1/ER-α. The tissue sections were scored for cytoplasmic and nuclear expression of GREB1 and nuclear expression of ESR1/ER-α in tumor cells and background ovarian stroma. The percentage of reactive tumor/stromal cells and intensity of reactivity (0, no staining; 1+ weak staining; 2+ moderately intense staining; and 3+ strong staining) was recorded.

## Results

### Patient Demographics

There were 115 patients with tissue from OM (n=6), PM (n=4), and primary CRC (n=105). The average age was 58 years old and most patients were male (57.4%), Caucasian (89.5%), and had microsatellite stable disease (69.6%). Most patients underwent cytotoxic chemotherapy (67%), while fewer underwent targeted therapy (38.3%) or radiation therapy (27.8%). The median overall survival for the primary CRC, OM, and PM populations was 7.1, 3.4, and 4.0 months, respectively.

### Gene Expression of Ovarian Metastases versus Primary Colorectal Cancer

Differential expression analysis was performed to compare OM (n=6) against primary CRC (n=105) tissue samples (Figure 1 and Figure 2a). There were 11,976 genes expressed by both OM and primary CRC, but only 83 (0.7%) of the genes were found to be differentially expressed. Of these 83 genes, 51 (61.4%) were upregulated and 32 (38.6%) were downregulated in OM (Table 1). Among upregulated genes, *PEG3, C7*, and *GREB1* had the highest differential expression in OM compared to primary CRC (all p<0.001). Among downregulated genes, *IGHG1, IGKV4-1*, and *MMP12* had the greatest differential expression in OM compared to primary CRC (all p<0.001).

**Figure 1.**
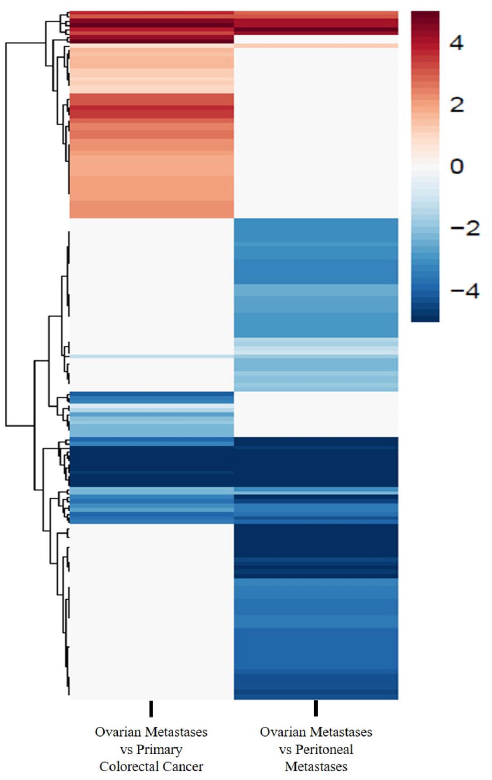
Heat map demonstrating gene expression profiling of ovarian metastases compared to primary colorectal cancer and peritoneal metastases.

**Figure 2.**
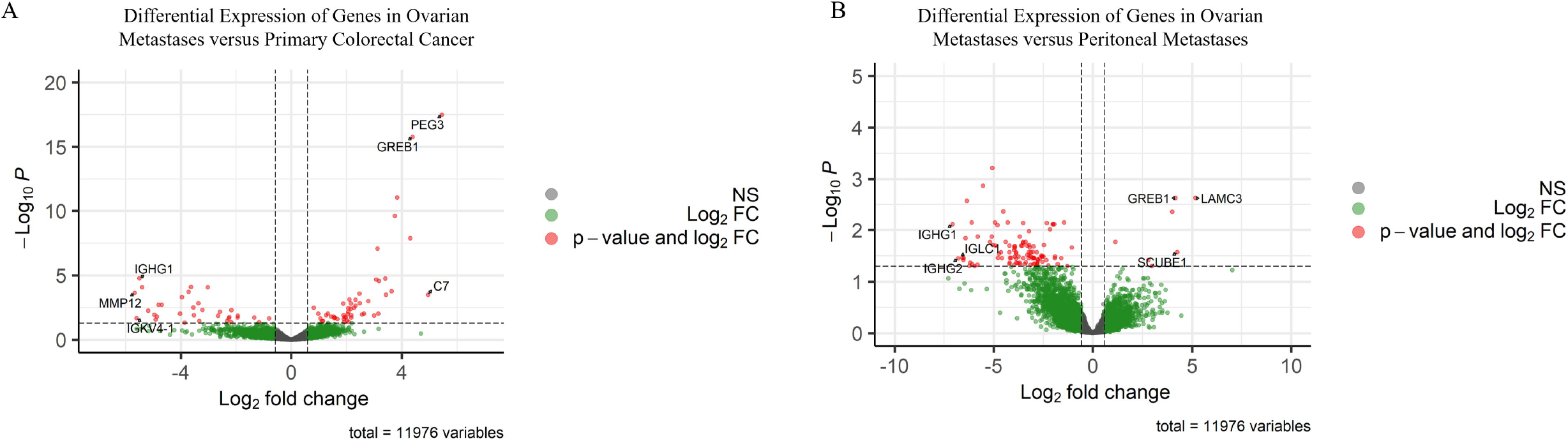
Volcano plot of differential expression of genes in A) ovarian metastases versus primary colorectal cancer samples and B) ovarian metastases versus peritoneal metastases samples

### Gene Expression of Ovarian Metastases versus Peritoneal Metastases

Differential expression analysis was performed to compare OM (n=6) against PM (n=4) tissue samples (Figure 1 and Figure 2b). There were 11,976 genes expressed by both OM and PM, but only 112 (0.9%) of the genes were found to be differentially expressed. Of these 112 genes, 7 (6.3%) were upregulated and 105 (93.7%) were downregulated in OM (Table 2). Among upregulated genes, *LAMC3, SCUBE1*, and *GREB1* had the highest differential expression in OM compared to PM (all p<0.001). Among downregulated genes, *IGLC1, IGHG2*, and *IGHG1* had the greatest differential expression in OM compared to PM (all p<0.001). Notably, *GREB1* was upregulated in OM compared to both PM and primary CRC.

### Low versus High GREB1 Expression - Survival Analysis

GEPIA2 is a web-based program that analyzes RNA sequencing expression data of 9,736 tumors and 8,587 normal samples from the Total Cancer Gene Atlas (TCGA) Program and Genotype-Tissue Expression (GTEx) Project.[11] With this program, overall survival and disease-free survival were compared for patients with high versus low *GREB1* expression in colon cancer samples. Low *GREB1* expression was associated with improved overall survival (p=0.029, Figure 3a) and disease-free survival (p=0.033, Figure 3b) in patients with colon cancer.

**Figure 3.**
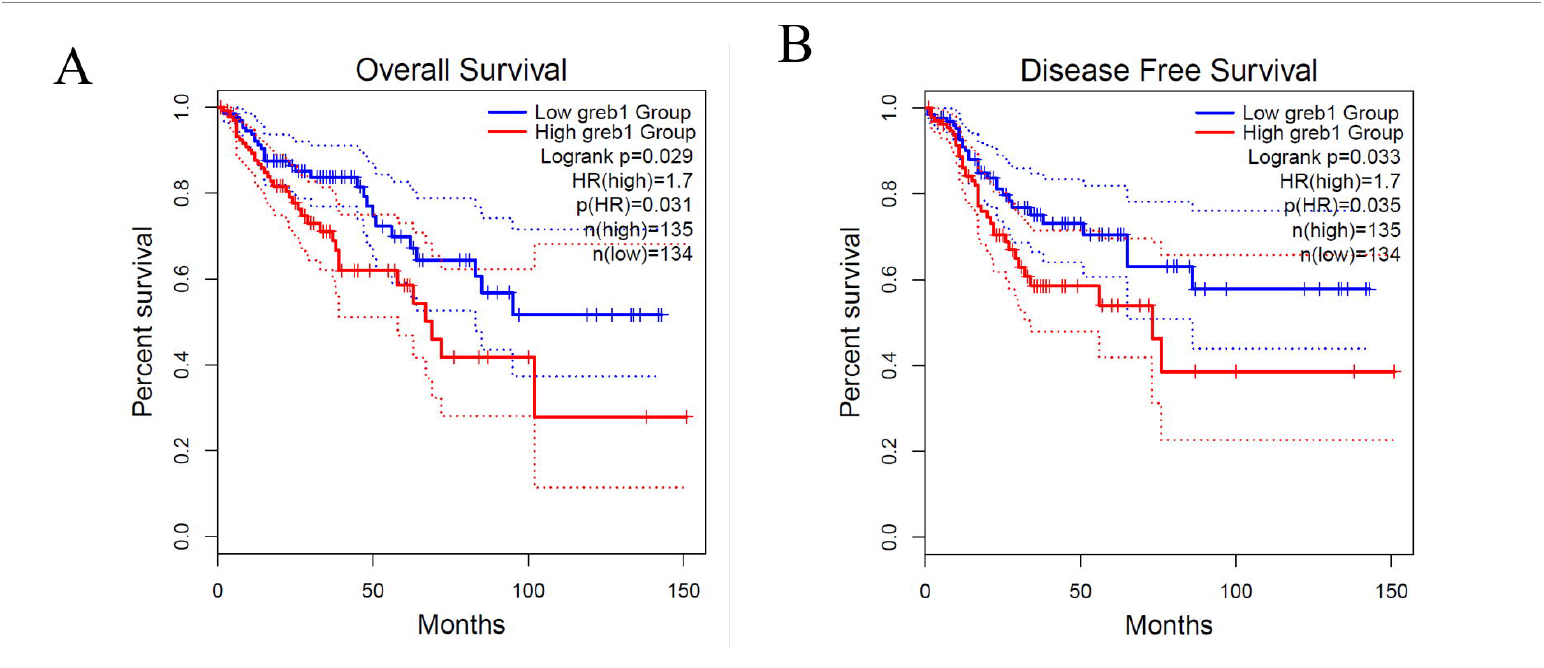
A) A) Overall survival and B) Disease free survival for low versus high GREB1 expression in colon cancer samples using the GEPIA2 web-based program.

### Analysis of Promoter Region of GREB1

There are two specific sites that ESR1 binds to ERE in the promoter region of *GREB1* (p=1.77e^-6^ and p=3.96e^-06^, respectively). As a result, there is a significant probability of direct binding of ESR1 to promoter regions of *GREB1,* and that ESR1 impacts the transcription of GREB1.

### Immunohistochemistry Staining for ESR1 and GREB1

IHC staining for ESR1 and GREB1 was performed on 11 patients with ovarian metastases from CRC. All 11 patients demonstrated nuclear staining of ESR1 only in the nearby ovarian stroma and not in the metastatic tumor (Figure 4a). GREB1 nuclear staining was positive in ovarian stroma for all 11 patients (Figure 4b). In the metastatic ovarian tumor, 7 patients (63.6%) demonstrated positive nuclear GREB1 staining and 5 patients (45.5%) showed positive cytoplasmic GREB1 staining in the ovarian metastasis (Figure 4c). This totals 8 of the 11 patients (72.7%). This suggests that GREB1 may be transcribed in the ovarian metastases independent of ERα signaling.

**Figure 4.**
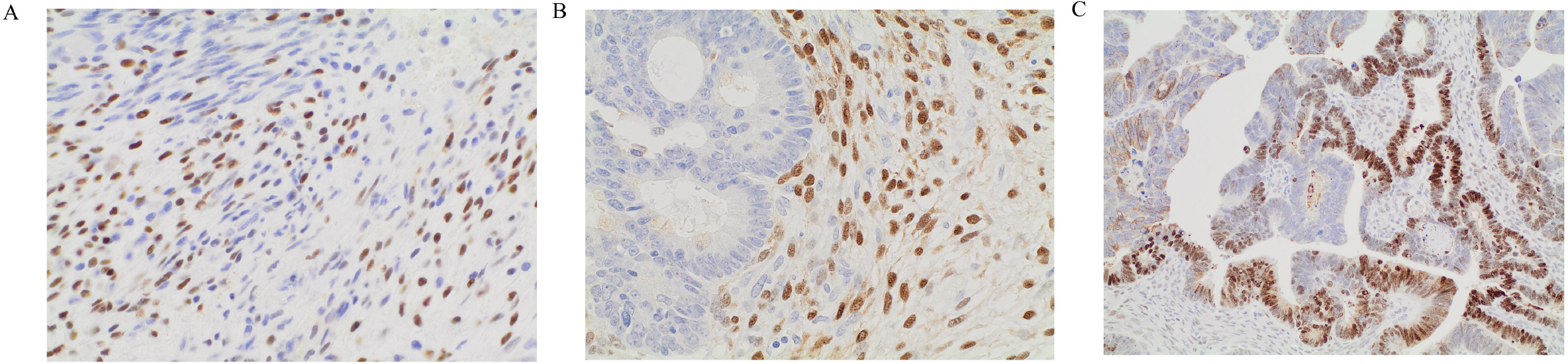
Representative images of A) ESRI positive immunohistochemistry nuclear staining in ovarian stroma and negative in ovarian metastasis B) GREB1 positive immunohistochemistry nuclear staining in ovarian stroma C) GREB1 positive immunohistochemistry nuclear and cytoplasmic staining in ovarian metastasis

## Discussion

This study is the first to compare the gene expression profile between primary CRC, PM, and OM. Transcriptomic analysis demonstrates a clear molecular distinction between OM, PM, and primary CRC. During this exploratory analysis, we found that *GREB1* was among the highest upregulated genes in OM compared to both PM and primary CRC. Furthermore, both ERE and ESR1 bind the upstream promoter region of *GREB1* at two separate sites, suggesting that transcription of *GREB1* is mediated by ESR1. On IHC, GREB1 and ESR1 expression was found in 100% of ovarian stroma, but only GREB1 expression was seen in the OM. This suggests that *GREB1* may be transcribed independently of ERα in OM and other mechanisms are driving its potential oncogenic effects. GREB1 is a potential novel target in the treatment of OM in CRC.

Genetic and molecular subtyping of metastatic CRC (mCRC) serves as the foundation for targeted therapy. DNA sequencing provides a static picture of the cell, whereas RNA sequencing provides a functional picture of the cell. Paired analysis of primary and metastatic tumors consistently shows concordance between genetic mutations.[12-18] However, this concordance among driver gene mutations does not extend to downstream gene expression.[19-21] The route of spread and unique microenvironment of different metastatic sites may select for a specific population of circulating cancer cells. In turn, cancer cells and native cells interact to establish the TME and impact cancer cell survival. The details of this process are still largely unknown.

The mechanism of metastasis is unclear for both OM and PM. Furthermore, the grouping of OM with PM as a common metastatic site is controversial given their different biologic behavior. Primarily, OM do not respond well to chemotherapy and confer a worse overall prognosis.[22-24] Our exploratory analysis found a clear difference in gene expression profiles between OM and PM. More specifically, *GREB1,* an early response gene in the estrogen receptor regulated pathway, was upregulated in OM. GREB1 is primarily shown to act as a regulator of hormone dependent growth in breast and prostate cancers.[25] However, one study showed that in CRC cell lines, overexpression of *GREB1* promoted cell proliferation and knockdown of *GREB1* resulted in reduced tumorigenesis.[26]

Nuclear receptors are transcription factors that regulate genes that control cell growth, homeostasis, differentiation, and metabolism. They are classified into three groups: hormone-mediated, metabolic, and orphan receptors. Dysregulation of these receptors can contribute to tumorigenesis.[27] ERβ (coded by *ESR2*) is the predominant estrogen receptor in normal colon mucosa. In previous studies, CRC specimens have been shown to present the loss or reduction of ERβ expression.[28] Low ERβ expression and high ERα (coded by *ESR1*) expression in these samples is associated with worse prognosis.[28, 29] This further demonstrates the significance of our data and the role that GREB1 may play in the development and progression of OM.

What is unclear based on our data is whether GREB1 expression is dependent on ERα in OM. The IHC staining showed that ovarian stroma stained positive for both GREB1 and ESR1, but that OM only stained positive for GREB1 (in both the cytoplasm and nucleus). This suggests that transcription of GREB1 in tumor cells may be activated independently of ERα. To support this, Matsumoto et al found that GREB1 is a target gene of Wnt/β-catenin signaling in hepatoblastoma formation.[30] Furthermore, GREB1 binds Smad2/3 in the nucleus and inhibits TGFβ signaling leading to cell proliferation in this mechanistic model. This same group recently showed that Wnt signaling stimulated GREB1 and HNF4α promoting hepatocellular carcinoma cell proliferation.[31]

There are limitations in this study that should be addressed. First, the sample size for OM and PM is small. Although despite the low sample size, statistical significance was still achieved. Second, bulk transcriptomic sequencing represents an overview of the gene expression of both cancer cells and the TME. Further research with spatial transcriptomics and single cell sequencing could reliably identify whether the upregulation of *GREB1* is originating from the cancer cells or the native epithelial cells of the ovary. This would also further clarify whether ERα is driving the transcription of *GREB1* in the OM.

Transcriptomic analysis clearly demonstrates molecular distinction between OM, PM, and primary CRC tissue. There was significant upregulation of *GREB1* in OM compared to PM and primary CRC. Additionally, *GREB1* contains two sites in the upstream promoter region that bind ERE and ESR1, suggesting transcription regulation mediated by ESR1. Tamoxifen, a selective estrogen receptor modulator, may be effective at overcoming drug resistance or inhibiting growth in CRC based on early *in vitro* and *in vivo* studies.[32] An estrogen receptor antagonist could serve as a novel targeted therapy or chemosensitizer for patients with OM or prevent OM in female patients with PM. However, the IHC data suggests that *GREB1* may also be activated independently of ERα. As such, further research with pre-clinical models to better delineate the mechanism of *GREB1* transcription and the downstream effects in CRC is necessary.

## Notes

The authors declare no potential conflicts of interest.

### Competing Interest Statement

The authors have declared no competing interest.

## References

1. Siegel, R.L., et al., Cancer statistics, 2023. CA: A Cancer Journal for Clinicians, 2023. 73(1): p. 17–48.

2. Testa, U., G. Castelli, and E. Pelosi, Genetic Alterations of Metastatic Colorectal Cancer. Biomedicines, 2020. 8(10).

3. Kajiyama, H., et al., Epidemiological overview of metastatic ovarian carcinoma: long-term experience of TOTSG database. Nagoya J Med Sci, 2019. 81(2): p. 193–198.

4. Segelman, J., et al., Epidemiology and prognosis of ovarian metastases in colorectal cancer. Br J Surg, 2010. 97(11): p. 1704–9.

5. Ganesh, K., et al., Clinical and genetic determinants of ovarian metastases from colorectal cancer. Cancer, 2017. 123(7): p. 1134–1143.

6. Wagstaff, J.F.R., et al., Perioperative outcomes of cytoreductive surgery for tumours of colorectal or appendiceal origin with ovarian involvement. J Surg Oncol, 2023. 128(1): p. 66–74.

7. Sekine, K., et al., Retrospective Analyses of Systemic Chemotherapy and Cytoreductive Surgery for Patients with Ovarian Metastases from Colorectal Cancer: A Single-Center Experience. Oncology, 2018. 95(4): p. 220–228.

8. Ritchie, M.E., et al., limma powers differential expression analyses for RNA-sequencing and microarray studies. Nucleic Acids Res, 2015. 43(7): p. e47.

9. Ikeda, K., K. Horie-Inoue, and S. Inoue, Identification of estrogen-responsive genes based on the DNA binding properties of estrogen receptors using high-throughput sequencing technology. Acta Pharmacologica Sinica, 2015. 36(1): p. 24–31.

10. Driscoll, M.D., et al., Sequence requirements for estrogen receptor binding to estrogen response elements. J Biol Chem, 1998. 273(45): p. 29321–30.

11. Tang, Z., et al., GEPIA2: an enhanced web server for large-scale expression profiling and interactive analysis. Nucleic Acids Research, 2019. 47(W1): p. W556–W560.

12. Kim, R., et al., Co-evolution of somatic variation in primary and metastatic colorectal cancer may expand biopsy indications in the molecular era. PLoS One, 2015. 10(5): p. e0126670.

13. Mogensen, M.B., et al., Genomic alterations accompanying tumour evolution in colorectal cancer: tracking the differences between primary tumours and synchronous liver metastases by whole-exome sequencing. BMC Cancer, 2018. 18(1): p. 752.

14. Sveen, A., et al., Intra-patient Inter-metastatic Genetic Heterogeneity in Colorectal Cancer as a Key Determinant of Survival after Curative Liver Resection. PLoS Genet, 2016. 12(7): p. e1006225.

15. Brannon, A.R., et al., Comparative sequencing analysis reveals high genomic concordance between matched primary and metastatic colorectal cancer lesions. Genome Biol, 2014. 15(8): p. 454.

16. Artale, S., et al., Mutations of KRAS and BRAF in primary and matched metastatic sites of colorectal cancer. J Clin Oncol, 2008. 26(25): p. 4217–9.

17. Fujiyoshi, K., et al., High concordance rate of KRAS/BRAF mutations and MSI-H between primary colorectal cancer and corresponding metastases. Oncol Rep, 2017. 37(2): p. 785–792.

18. Bhullar, D.S., et al., Biomarker concordance between primary colorectal cancer and its metastases. EBioMedicine, 2019. 40: p. 363–374.

19. Kamal, Y., et al., Transcriptomic Differences between Primary Colorectal Adenocarcinomas and Distant Metastases Reveal Metastatic Colorectal Cancer Subtypes. Cancer Res, 2019. 79(16): p. 4227–4241.

20. Lenos, K.J., et al., Molecular characterization of colorectal cancer related peritoneal metastatic disease. Nat Commun, 2022. 13(1): p. 4443.

21. Yaeger, R., et al., Clinical Sequencing Defines the Genomic Landscape of Metastatic Colorectal Cancer. Cancer Cell, 2018. 33(1): p. 125-136.e3.

22. Mori, Y., et al., Clinical outcomes of women with ovarian metastases of colorectal cancer treated with oophorectomy with respect to their somatic mutation profiles. Oncotarget, 2018. 9(23): p. 16477–16488.

23. Lee, S.J., et al., Survival benefit from ovarian metastatectomy in colorectal cancer patients with ovarian metastasis: a retrospective analysis. Cancer Chemother Pharmacol, 2010. 66(2): p. 229–35.

24. Goéré, D., et al., The differential response to chemotherapy of ovarian metastases from colorectal carcinoma. Eur J Surg Oncol, 2008. 34(12): p. 1335–9.

25. Hodgkinson, K., et al., GREB1 is an estrogen receptor-regulated tumour promoter that is frequently expressed in ovarian cancer. Oncogene, 2018. 37(44): p. 5873–5886.

26. Kochi, M., et al., Oncogenic mutation in RAS-RAF axis leads to increased expression of GREB1, resulting in tumor proliferation in colorectal cancer. Cancer Sci, 2020. 111(10): p. 3540–3549.

27. Triki, M., et al., Expression and role of nuclear receptor coregulators in colorectal cancer. World J Gastroenterol, 2017. 23(25): p. 4480–4490.

28. Topi, G., et al., Association of the oestrogen receptor beta with hormone status and prognosis in a cohort of female patients with colorectal cancer. Eur J Cancer, 2017. 83: p. 279–289.

29. Topi, G., et al., Combined Estrogen Alpha and Beta Receptor Expression Has a Prognostic Significance for Colorectal Cancer Patients. Front Med (Lausanne), 2022. 9: p. 739620.

30. Matsumoto, S., et al., GREB1 induced by Wnt signaling promotes development of hepatoblastoma by suppressing TGFβ signaling. Nat Commun, 2019. 10(1): p. 3882.

31. Matsumoto, S., et al., Wnt Signaling Stimulates Cooperation between GREB1 and HNF4α to Promote Proliferation in Hepatocellular Carcinoma. Cancer Res, 2023. 83(14): p. 2312–2327.

32. Morad, S.A.F., et al., Tamoxifen magnifies therapeutic impact of ceramide in human colorectal cancer cells independent of p53. Biochemical Pharmacology, 2013. 85(8): p. 1057–1065.

